# BacilloFlex: A modular DNA assembly toolkit for *Bacillus subtilis* synthetic biology

**DOI:** 10.1101/185108

**Authors:** Niels Wicke, David Radford, Valeria Verrone, Anil Wipat, Christopher E. French

## Abstract

*Bacillus subtilis* is a valuable industrial production platform for proteins, a bacterial model for cellular differentiation and its endospores have been proposed as a vehicle for vaccine delivery. As such *B. subtilis* is a major synthetic biology chassis but, unlike *Escherichia coli*, lacks a standardized toolbox for genetic manipulation. EcoFlex is a versatile modular DNA assembly toolkit for *E. coli* synthetic biology based on Golden Gate cloning. Here we introduce BacilloFlex, an extension of the EcoFlex assembly standard to *B. subtilis*. Transcription units flanked by sequences homologous to loci in the *B. subtilis* genome were rapidly assembled by the EcoFlex standard and subsequently chromosomally integrated. At present, BacilloFlex includes a range of multi-functional *B. subtilis* specific parts with applications including metabolic engineering, biosensors and spore surface display. We hope this work will form the foundation of a widely adopted cloning standard for *B. subtilis* facilitating collaboration and the sharing of parts.

## 1. Introduction

Since the publication of its genome in 1997 (Kunst et al., 1997), the naturally competent gram-positive spore-forming bacterium *B. subtilis* has attracted considerable academic and industrial interest. Arguably, the amenability of *B. subtilis* to genetic modification complemented by further useful properties, like biosafety and capacity for protein production, has led to its rise as the world’s second best studied bacterium after *Escherichia coli*.

Unlike *E. coli, B. subtilis* lacks toxins or immunogenic lipopolysaccharides. In fact, *B. subtilis* is designated Generally Regarded as Safe (GRAS) by the U.S. Food and Drug Administration, an important economic and regulatory detail for bioprocesses (Schallmey et al., 2004). Furthermore, (heterologous) protein production in gram-positives is relatively simple because polypeptides tagged with an N-terminal signal sequence are directly secreted into the extracellular medium (rather than the periplasm) via the Sec pathway, and secreted yields readily reach 20-25 g/L (van Dijl and Hecker, 2013, Harwood and Cranenburgh, 2008). The fact that 900 tons of native proteases for detergents are produced using *Bacillus* spp. per year in Europe is testament to their utility for protein production (van Dijl and Hecker, 2013). Recombinant *B. subtilis* is additionally a production platform for compounds with applications in the food, pharmaceutical and biofuel industries, such as riboflavin and (R,R)-2,3-butanediol (Hao et al., 2013).

Sporulation in *B. subtilis* has also attracted academic interest as a model for cellular differentiation. The study of *B. subtilis* and its sporulation process offers important design principles for gene network architecture and molecular mechanisms of cell differentiation (Narula et al., 2016).

Described first by Spizizen (1958) competence in *B. subtilis* is a post-exponential phase, quorum-signalling regulated, physiological state where 10-20% of cells upregulate machinery for the uptake and chromosomal integration of extracellular double-stranded DNA (Dubnau, 1991). Double-stranded DNA is nicked and actively internalized in a single-stranded form where it is coated by RecA mediating homologous recombination with the host chromosome (Yadav et al., 2014). The competence network is conserved across many *Bacillus* spp. but activation thereof is often laborious, for example by overexpression of master regulator ComK. In *B. subtilis*, by contrast, -80°C stocks with reproducible levels of competence can be prepared with ease by growth in a defined medium and timely harvesting of cells (Bron, 1990).

Finally, relatives of *B. subtilis* include *Bacillus anthracis* the etiological agent of anthrax (Mock and Fouet, 2001), *Bacillus amyloliquefaciens*, a rhizobacterium used in agriculture (Chen et al., 2016), and the *Clostridium* genus featuring the medically important *Clostridium perfringens* and *Clostridium botulinum*, and *Clostridium acetobutylicum* used in biofuel production (Chen et al., 2016, Mock and Fouet, 2001, Lee et al., 2008). Knowledge gathered in *B. subtilis*, rather than *E. coli*, is likely to translate to well to said organisms.

Taken together, *B. subtilis* is of academic and industrial value, a model platform organism for related important gram-positives, and amenable to genetic modification, making it an excellent synthetic biology chassis. Despite the evident importance of *B. subtilis* a lack of publicly accessible, flexible and standardized genetic tools hinders research (Radeck et al., 2013, Guiziou et al., 2016).

Golden Gate (GG) cloning utilizes the properties of Type IIS restriction enzymes (T2SRE) to enable rapid combinatorial assembly of parts in a pre-defined order with high fidelity and efficiency in a single tube (Engler et al., 2008). EcoFlex, developed by Moore et al. (2016), places GG technology in a standardized hierarchical assembly scheme. A first GG reaction assembles bio-parts into a TU, which in turn are combined into a multi-TU construct by a second GG reaction. Moore et al. (2016) demonstrate the power of EcoFlex by assembling a 20-gene (68-part) containing 31 kb plasmid. Providing functionality, the EcoFlex toolkit includes a range of characterized parts such as promoters, RBS and terminators with variable strengths, N-terminal tags and reporters characterized *in vitro* and *in vivo*. Characterization of each re-usable part or module facilitates the bottom-up approach of rationally designing gene networks, where *in silico* modelling precedes assembly and *in vivo* testing, thereby bypassing laborious trial-and-error projects (Radeck et al., 2013, Rebatchouk et al., 1996).

Here we present BacilloFlex an extension of the EcoFlex assembly standard and toolkit. BacilloFlex adopts the EcoFlex work-flow to assemble an integrative plasmid composed of a pair of homology arms mediating chromosomal integration, an antibiotic resistance marker for selection, and a user-defined TU (Figure 1). The BacilloFlex toolkit comprises a set of *B. subtilis* specific parts, including eight promoters, conforming to and compatible with the EcoFlex assembly standard. Taken together, BacilloFlex combines the ease of genetic modification of naturally competent *B. subtilis* with the power of DNA assembly endowed by EcoFlex.

**Figure 1.**
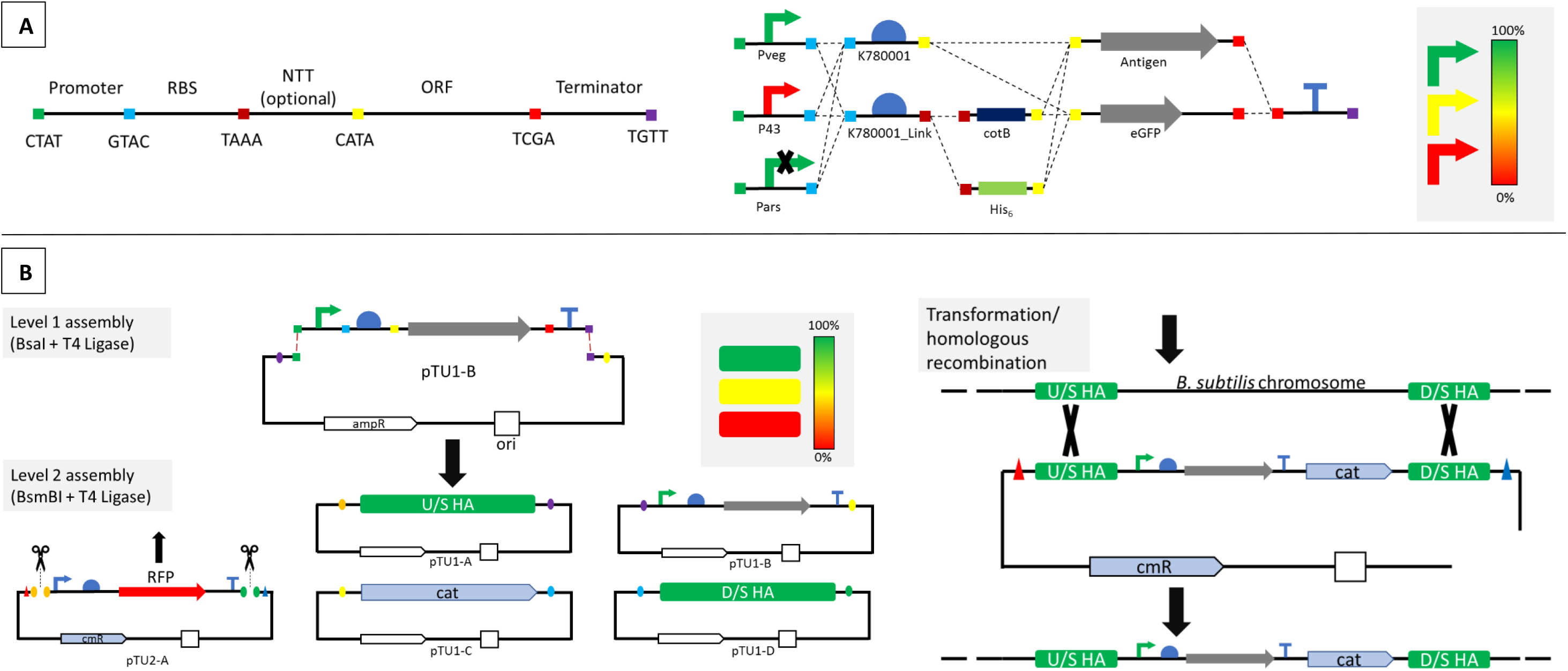
BacilloFlex overview. **(A) EcoFlex level 0 design** Bio-parts (promoter, RBS (ribosome binding site), NTT (N-terminal tag), ORF (open reading frame), Terminator) are designated level 0. The 4 base fusion sites (overhangs generated by T2SRE) (coloured squares) of each part class are detailed below the black line. Note the downstream RBS fusion site may be TAAA or CATA for including or excluding an N-terminal tag respectively. The flexibility in combinatorial assembly of level 1 constructs or transcription units (accepting vector not shown) from level 0 parts is illustrated. For example, analogous TUs with promoters of differing strengths, for fine-tuning multi-gene pathways, are easily assembled. **(B) Assembly and integration workflow.** Level 0 parts are assembled into pTU1-B according to the EcoFlex standard. Level 1 plasmids are principally maintained in *E. coli*. The level 2 assembly includes the user-defined TU (pTU1-B), a pair of ‘fake’ level 1 TU homology arms (pTU1-A/D) and a chloramphenicol resistance cassette (cat)(pTU1-C). The resulting level 2 construct thus flanks a user defined TU and cat, enabling selection, with upstream (U/S) and downstream (D/S) homology arms (HA) that mediate chromosomal integration by homologous recombination. HAs integrating at loci with different properties, such as high (green) or low (red) expression levels, are available (see Figure 2). Figure inspired by Moore et al. (2016).

## 2. Results & Discussion

### 2.1 Construction of parts

To begin building a library of *B. subtilis* specific parts a range of level 0 parts were constructed, including promoters with variable predicted properties (Table 1). In addition, five pairs of homology arms and a chloramphenicol resistance cassette were assembled (Table 2). The choice of homology arms was inspired by the work of Sauer et al. (2016) (Figure 2). Promoters, NTTs, RBS and Xer-cise parts were cloned into pBP-lacZ (EcoFlex #72948), and ORFs into pBP-ORF (EcoFlex #72949). Upstream homology arms, downstream homology arms and antibiotic resistance cassettes were cloned into pTU1-A-RFP (EcoFlex #72935), pTU1-D-RFP (EcoFlex #72938) and pTU1-C-RFP (EcoFlex #72937) respectively. Correct cloning was tested by colony polymerase chain reaction (PCR) and confirmed by complete sequencing (see Table S1 for primers). See the Materials and Methods section for a more detailed account of part construction.

**Table 1.** Level 0 parts.

| Type of part | Name | Description | Relevant references |
| --- | --- | --- | --- |
| Promoter | $P_{43}$ | BBa_K143013, strong activity during exponential and stationary phase | Wu et al. (1991) |
| Promoter | $P_{veg}$ | BBa_K823003, strong promoter active during sporulation and vegetative growth | Fukushima et al. (2003) |
| Promoter | $P_{liaG}$ | BBa_K143012, weak constitutive promoter of *liaGFSR* operon involved in detection of cell wall damage | Jordan et al. (2006) |
| Promoter | $P_{spac}$ | Repressed by LacI (not natively produced in *B. subtilis* 168) | Sun et al. (2009), Liu et al. (2014) |
| Promoter | $P_{ars}$ | Promoter of *ars* operon. Repressed by ArsR, induced by arsenate or arsenite | Sato and Kobayashi (1998) |
| Promoter | $P_{cotB}$ | Predicted promoter of *cotB* | This study |
| Promoter | $P_{cotY}$ | Predicted promoter of *cotY* | This study |
| Promoter | $P_{cotZ}$ | Predicted promoter of *cotZ* | This study |
| RBS | K780001 | BBa_K780001, proposed strong RBS in *B. subtilis*, downstream fusion site to ORFs | n/a |
| RBS | K780001_Link | As K780001 but including downstream fusion site to N-terminal tags | n/a |
| N-terminal Tag | CotB | First 624 bp of *cotB* (outer spore coat protein) | This study |
| N-terminal Tag | CotY | *cotY* excluding the stop codon (spore crust protein) | This study |
| N-terminal Tag | Epr | Signal peptide of *epr* | Brockmeier et al. (2006) |
| N-terminal Tag | PhrK | Signal peptide of *PhrK* | Brockmeier et al. (2006) |
| ORF | XylE | BBa_K118021, encodes catechol-2,3-dioxygenase, from *Pseudomonas putida* TOL pWW0. | Polissi and Harayama (1993) |
| Xer-cise | XC5 | *B. subtilis dif* sequence with upstream CTAT and downstream TAAA fusion site | Bloor and Cranenburgh (2006) |
| Xer-cise | XC3 | *B. subtilis dif* sequence with upstream CATA and downstream TGTT fusion sites | Bloor and Cranenburgh (2006) |
| Xer-cise | catXC | Chloramphenicol resistance cassette (expressing chloramphenicol acetyltransferase) with upstream TAAA and downstream CATA fusion site | n/a |

For complete promoter and RBS sequences see Table S2.

**Table 2.**
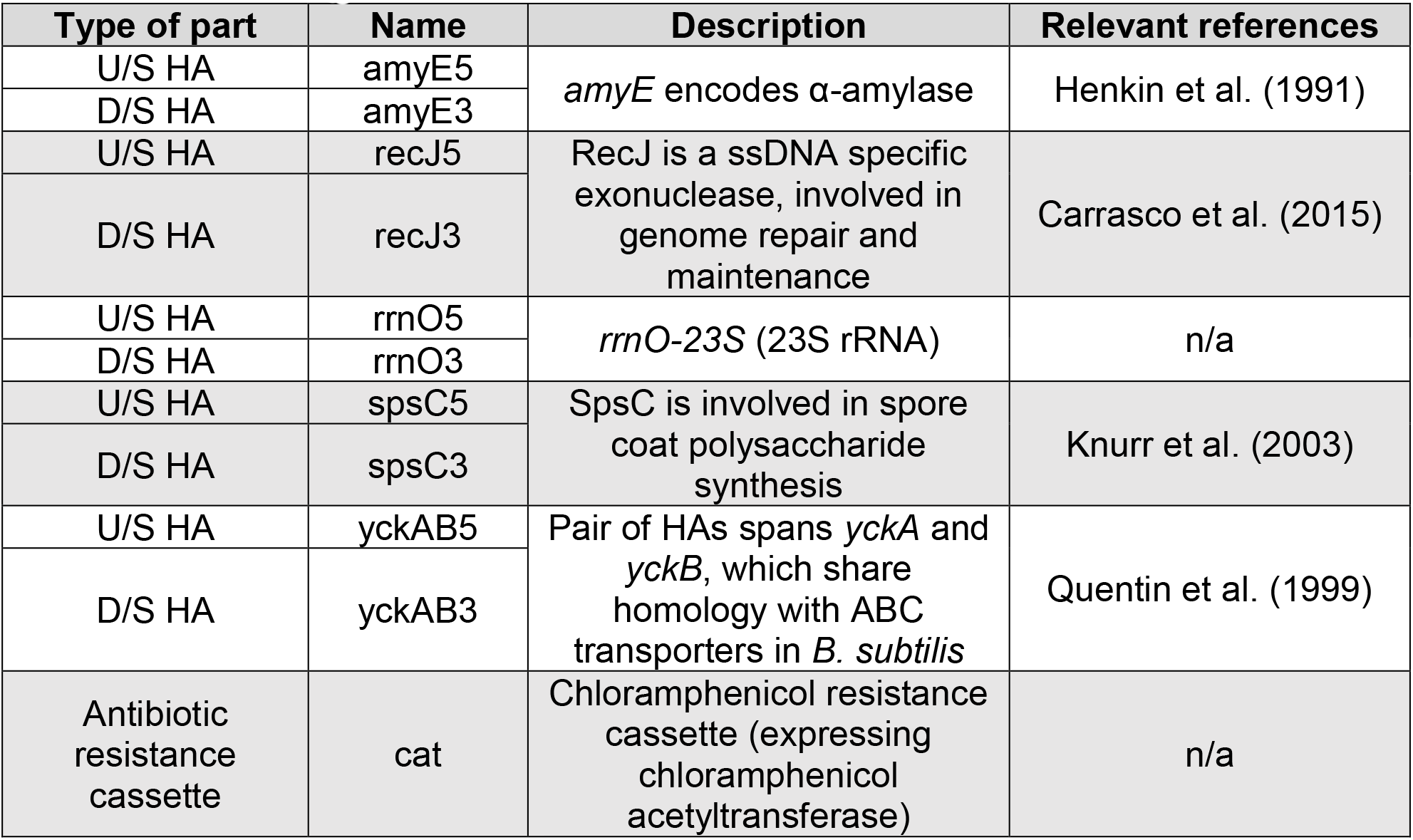
Level 1 parts.

**Figure 2.**
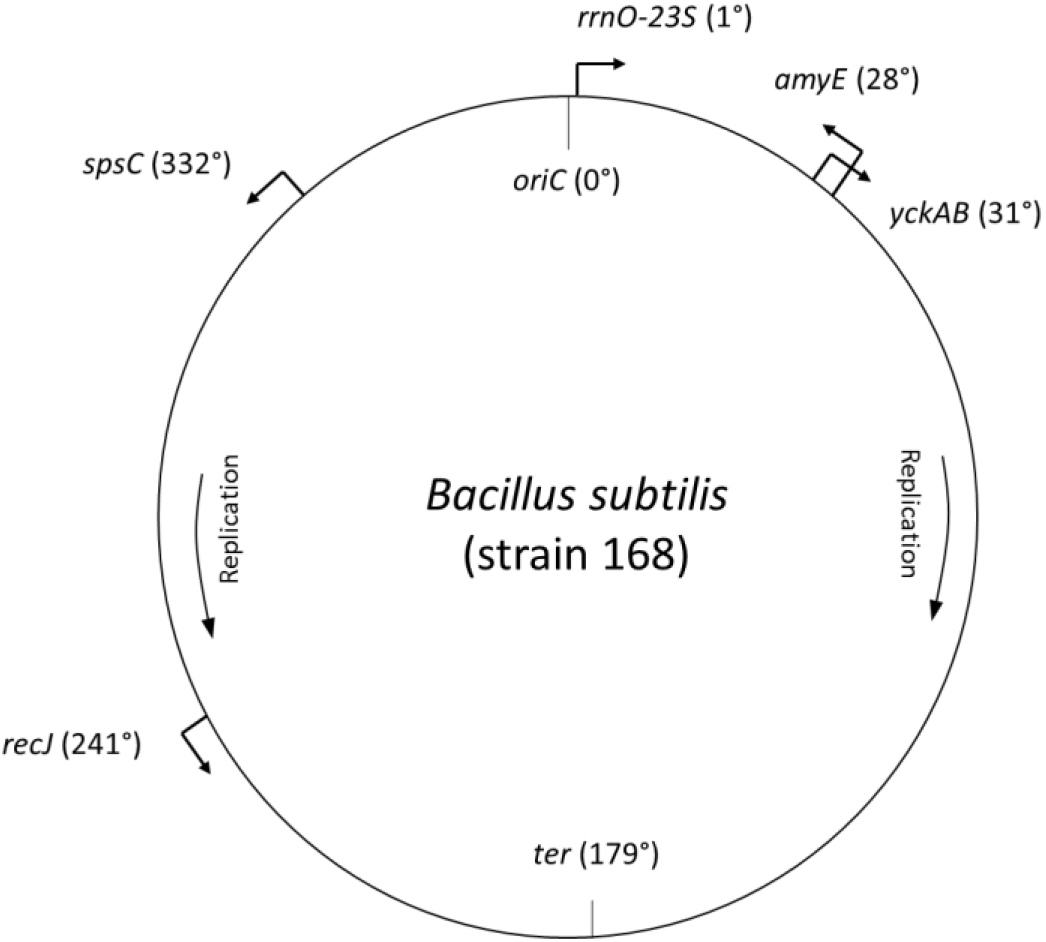
Map of integration loci on the *B. subtilis* genome. The gene dosage effect refers to the influence of genome position, specifically the proximity to the origin of replication, on the expression level of genes during rapid growth due to differences in copy number. Unbiased analysis by Sauer et al. (2016) demonstrated that the gene dosage effect in *B. subtilis* can account for a five-fold difference in expression levels and that integration into the loci above, amongst others, is not lethal. That is to say, the respective genes are nonessential. Arrows indicate the direction of the gene. 1° corresponds to approximately 11708 bp. The origin *(oriC)* and terminus *(ter)* of replication are marked. The bi-directional nature of chromosome replication is shown. Figure adapted from Sauer et al. (2016).

### 2.2 Assembly and integration

To test our proposed strategy for chromosomal integration we assembled a level 2 construct flanked by *amyE* HAs (pTU1-A-amyE5 + pTU1-D-amyE3) and carrying the chloramphenicol resistance cassette (pTU1-C-cat). Position 2 (pTU1-B) was occupied by one of four analogous TUs expressing enhanced green fluorescent protein (eGFP, EcoFlex #72960) under control of P*_43_*, P*_veg_*, P*_liaG_* or P*_ars_*. The RBS was BBa_K780001 and the terminator BBa_B0015 (EcoFlex #72998). The assembled integrative plasmid was transformed into *B. subtilis* and chromosomal insertion confirmed by colony PCR (Figure 3). An increase in size of ∼1.4 kb between wild-type and transformants is consistent with correct insertion of respective level 2 constructs by a double crossover.

**Figure 3.**
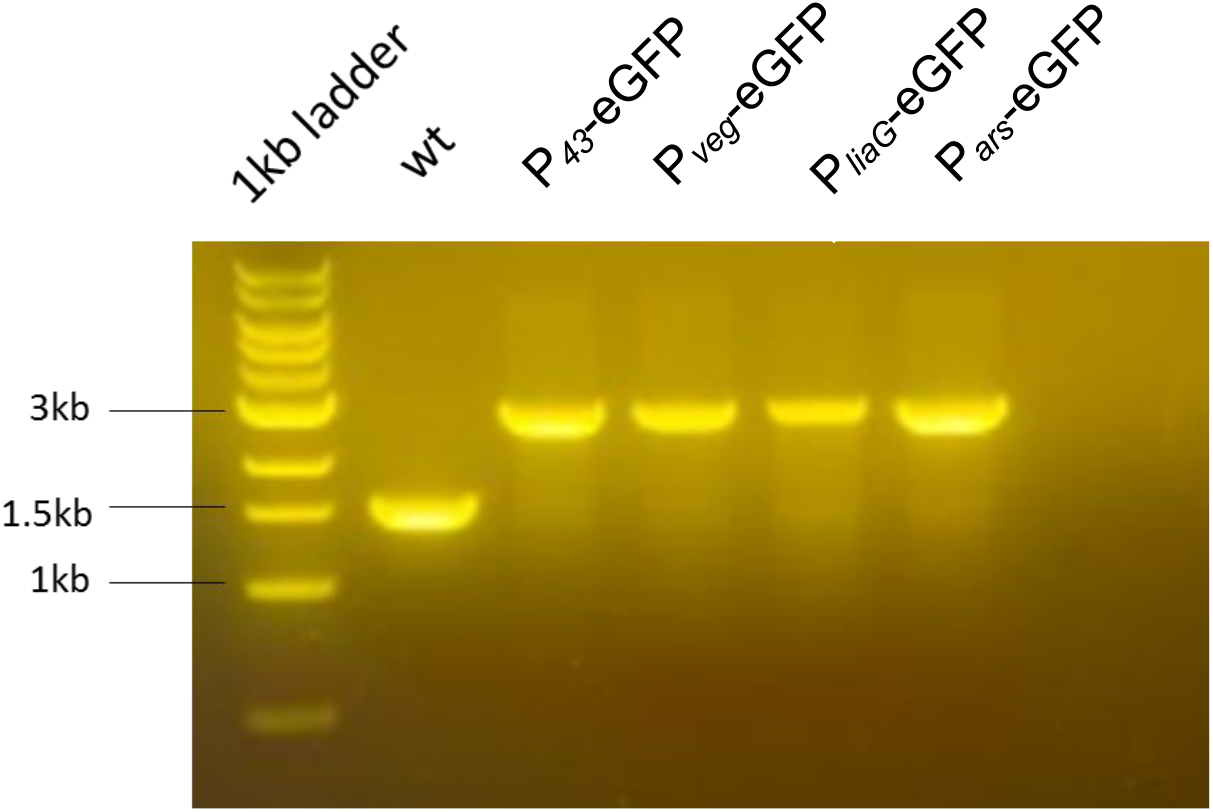
Construct integration at *amyE* locus. Colony PCRs with primers flanking *amyE* insertion locus (amyE_F, amyE_R) of *B. subtilis* transformed with integrative level 2 constructs. Wild-type (wt) *B. subtilis* included as size-marker. Expected wt band size was 1590 bp, and 2927 - 2992 bp for transformants depending on promoter size.

### 2.3 Promoter activity at different integration loci

Fine-tuning of synthetic multi-gene pathways requires control beyond maximal levels of transcription. Toward that end, and demonstrating the utility of BacilloFlex, we sought to quantify the activity of a set of BacilloFlex promoters at different integration loci. First, TUs expressing eGFP under control of different promoters (P*_43_*, P*_veg_*, P*_liaG_*, P*_ars_* and P*_spac_*) with K78001 and BBa_B0015 were assembled in pTU1-B. Next, all twenty-five combinations of TUs and HA pairs were constructed, including *cat* for selection, and transformed into *B. subtilis*. eGFP production during exponential growth phase was quantified to investigate the gene dosage effect and promoter activity (Figure 4). We observed a large spectrum of promoter activity with a 125-fold dynamic range (P*_43_ amyE* to P *_ars_ rrnO*).

### 2.4 Inducible promoters and arsenic biosensors

Inducible promoters provide tighter control over gene expression as required by many applications, for instance if the product is toxic. Our previous data (Figure 4) demonstrate that P*_spac_* and P*_ars_* are active. The activity of P*_spac_* in *B. subtilis* has been shown by others (Sun et al., 2009, Liu et al., 2014), although it is repressed by artificial overproduction of LacI which is relieved by addition of 5-bromo-4-chloro-3-indolyl-β- D-galactopyranoside (IPTG).

P*_ars_* is the cognate promoter of the arsenic resistance operon *(ars)* in *B. subtilis*. In the absence of arsenate or arsenite ArsR represses transcription of the *ars* operon (Sato and Kobayashi, 1998). To assess P*_ars_* inducibility, eGFP production by a *B. subtilis* strain expressing eGFP under control of P*_ars_* at the *amyE* locus in different concentrations of arsenate was quantified (Figure 5). There were no differences in P*_ars_* inducibility between stationary and exponential phase. The data reiterates that P*_ars_* is a leaky promoter but in this configuration its activity increases by 9 ± 8%, 40 ± 10%, 278 ± 18% and 440 ± 50% in the presence of 0.1, 1, 10 and 100 ppb (μg/L) arsenate (AsO_4_^3-^) respectively.

### 2.5 Additional parts

Offering additional functionality the BacilloFlex kit also includes the cognate secretion signals of *Epr* and *PhrK* (Brockmeier et al., 2006), an excisable chloramphenicol resistance marker by Xer-mediated recombination (Bloor and Cranenburgh, 2006), and the reporter gene *xylE* (Polissi and Harayama, 1993) (Table 1). These parts have been constructed and sequence confirmed. Their biological activity has not been tested in this study, but previously by others (see references). Similarly, sequence confirmed fragments of spore coat proteins CotB and CotY as N-terminal tags for spore surface display and their cognate promoters are available but have not been tested (Table 1, see supplementary information for sequence).

**Figure 4.**
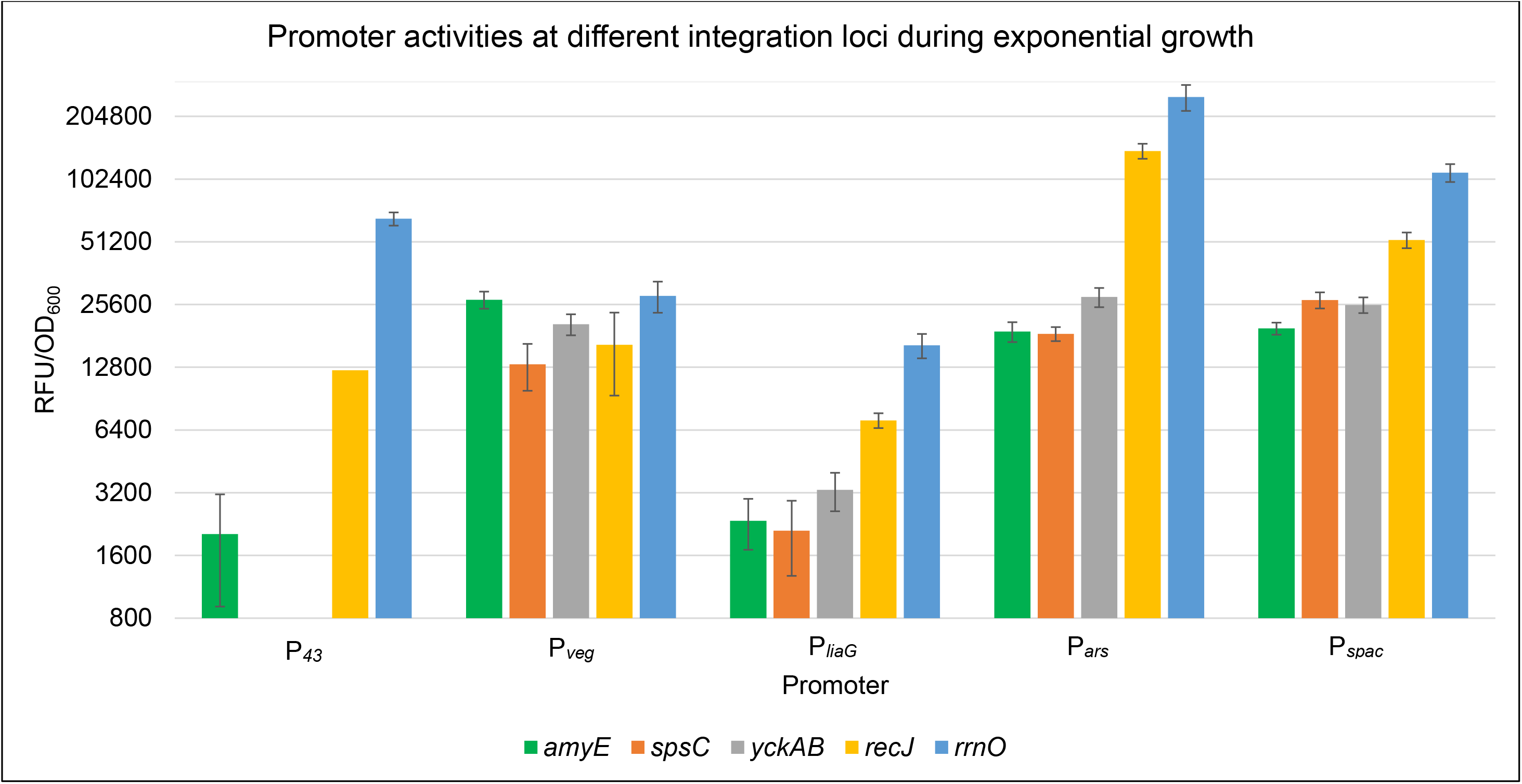
Promoter activities at different integration loci during exponential growth. Average relative fluorescence units (RFU) corrected for cell density (absorbance at 600 nm, OD_600_) of strains expressing eGFP, from different promoters at different genomic loci, during exponential growth over a 50 min period are shown. Error bars indicate standard deviation of three technical replicates. Wild-type *B. subtilis* RFU/OD_600_ has been subtracted.

**Figure 5.**
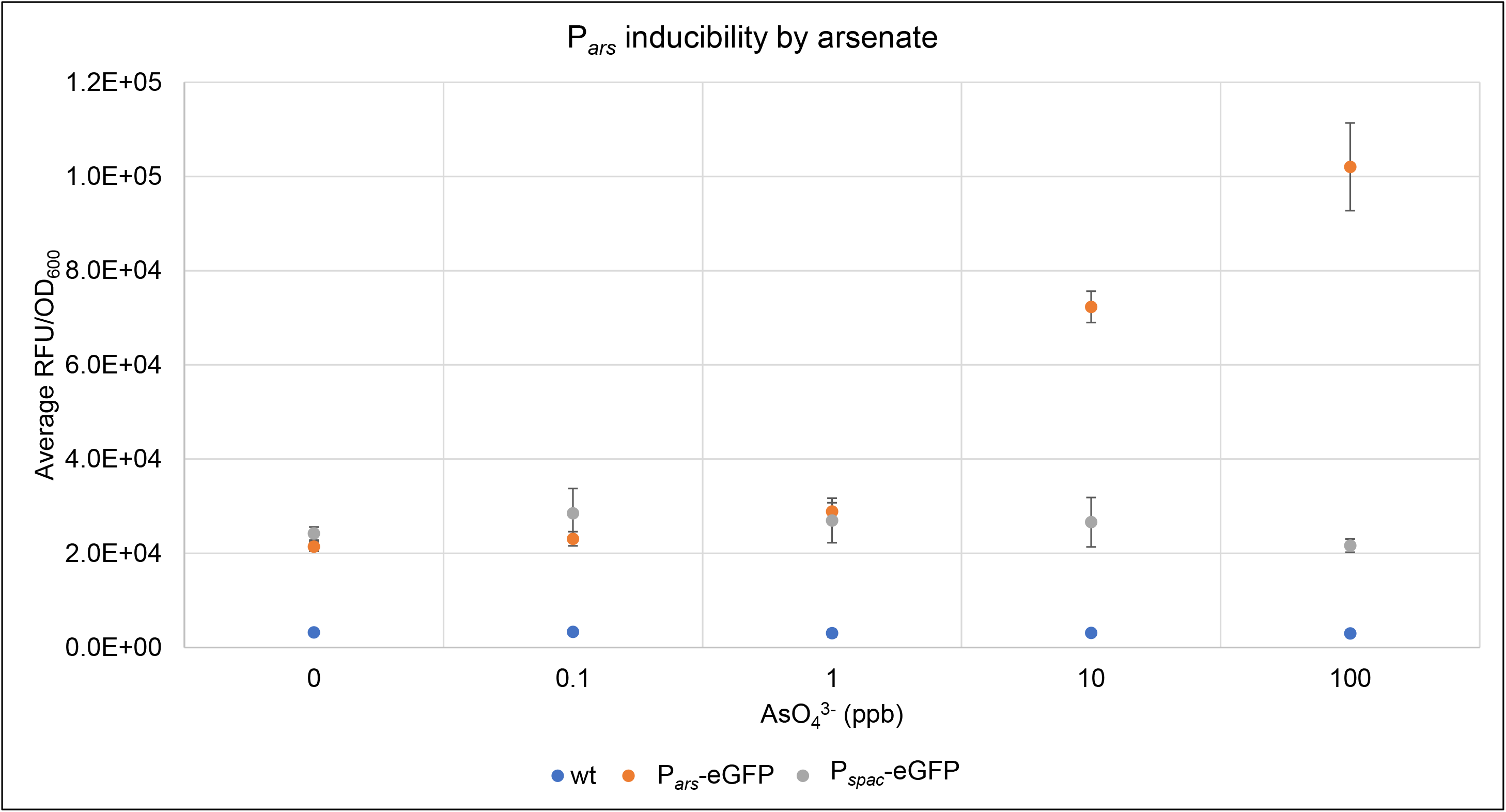
P*_ars_* inducibility during stationary phase. Average relative fluorescence units (RFU) corrected for cell density (OD_600_) of strains expressing eGFP during stationary phase in different concentrations of arsenate over a 60 min period are shown. Error bars indicate standard deviation of technical triplicates. 10 ppb or 10 μg/L arsenate (AsO_4_^3-^) contains 5.4 μg/L arsenic (As). P*_ars_*-eGFP and P*_spac_*-eGFP constructs were integrated at the *amyE* locus.

## 3. Discussion

We have demonstrated the feasibility of a modular DNA assembly toolkit, based on the EcoFlex standard (Moore et al., 2016), for chromosomally modifying *B. subtilis*. Combining the power of Golden Gate cloning, modularity and the natural competence of *B. subtilis*, TUs can be assembled from basic bio-parts and integrated into the host genome within five days.

It is worth noting that transforming level 2 Golden Gate reactions directly into *B. subtilis* routinely gave 0-5 colony forming units (CFUs) when 20% of transformation culture was plated onto selective media. We therefore recommend, to forego unnecessary repetition, transforming level 2 Golden Gate reactions into both *E. coli* and *B. subtilis*, or to use remaining *B. subtilis* transformation cultures as inoculum for overnight cultures (see Materials and Methods, *‘B. subtilis* transformation’). Preliminary tests suggest increasing DNA concentration 1.5-fold in level 2 assemblies and subsequent transformations raises CFU numbers. Snap-freezing aliquots when preparing *B. subtilis* competent cells may further improve transformation efficiency.

At present our BacilloFlex kit contains a range of basic parts, including promoters, pairs of homology arms, a reporter, secretion signals, and spore coat protein N-terminal tags providing functionality in a variety of applications including metabolic engineering.

The BacilloFlex promoter-integration locus combinations provide a spectrum of gene expression strengths (Figure 4); however, the range of P*_43_*, P*_liaG_*, P*_ars_* and P*_spac_* activity at different loci exceeds the 5-fold difference of the gene dosage effect described by Sauer et al. (2016), likely due to read-through transcription which was mitigated in that study by use of terminator flanked reporters. High expression levels at *recJ* and *rrnO* are unsurprising given that their products are a single-strand DNA-specific exonuclease involved in recombinational DNA repair and genome maintenance, and the 23S rRNA respectively. Disruption of *recJ* is reported to increase susceptibility to killing by DNA damaging agents (Carrasco et al., 2015). This should be considered if *recJ* is chosen as insertion locus for strains with applications where this may be problematic, for instance in long continuous fermentations. Overall, the results here broadly conform to the predicted activity of promoters and integration loci (Table 1 and 2, Figure 3), but are perhaps best taken for the functionality they provide. Definite conclusions regarding the activity of individual promoters and integration loci should be reserved for additional work including the testing of biological replicates and *in vitro* experiments.

The data in Figure 5 demonstrate the sensitivity and inducibility of P*_ars_*. The linear relationship, above a minimal concentration of inducer, between arsenate concentration and P*ars* activity observed here is consistent with previously published results of Sato and Kobayashi (1998) who also note that in *arsR* expressing constructs maximal P*_ars_* induction occurs at arsenate levels 1000-fold higher than tested here (10 μM or 10,000 ppb). The World Health Organization’s recommended maximum concentration for arsenic in drinking water is 10 μg/L (10 ppb As, 18.5 ppb AsO_4_^3-^) (Flanagan et al., 2012). Given that strong induction occurs below this threshold, arsenate may therefore be used as an inexpensive inducer in bioprocesses, although overproduction of ArsR is likely needed to obtain tighter control over expression which may affect sensitivity to a point where arsenate concentration become hazardous. The sensitivity of P*_ars_* to arsenate reiterates its utility in a biosensor configuration. We have since used BacilloFlex to construct a chromosomally integrated version of an arsenic biosensor previously developed in our lab (Wicke, Radford and French, manuscript in preparation).

Translational fusion of spore coat proteins, such as CotB or CotC, to antigens has been demonstrated as a potent vaccine configuration (Zhou et al., 2017, Amuguni and Tzipori, 2012, Pohl et al., 2013). The inherent resistance of *B. subtilis* spores to a variety of physical stresses and absence of nutrients contributes to their utility as an antigen delivery vehicle because it facilitates manufacture, transport and storage. In addition, they have been shown to act as an adjuvant (Cutting et al., 2009, de Souza et al., 2014). Therefore, future work should include testing the BacilloFlex spore coat fragments, perhaps toward developing a rapid-development vaccine platform where the efficiency of BacilloFlex enables quick assembly of recombinant antigen-presenting spores.

In addition, the assembly of level 3 integrative plasmids for insertion of multiple TUs should be tested. It is noted that the present design of EcoFlex precludes C-terminal tags, necessitates the inclusion of a CATATG string in N-terminal fusions and that palindromic fusion sites (e.g. promoter-RBS, GTAC) may reduce assembly efficiency.

While this modular cloning standard is still in its infancy these design features, if possible, should be reconsidered. Preliminary work in our lab has shown that “old” and “new” parts can be made retro-compatible through the inclusion of short oligo-linkers (Marcos Valenzuela, personal communication).

Finally, domestication of a replicative vector in *B. subtilis* to the EcoFlex standard is also desirable. A potential backbone is pTG262 (BBa_I742123), which is a broad host-range vector with a gram-positive origin of replication that is also compatible with *E. coli* (Shearman et al., 1989). It has been shown to replicate in *B. subtilis* and lactic acid bacteria including *Lactobacillus* spp. and *Lactococcus* spp. Hence, it may prove particularly valuable as it will extend the EcoFlex standard to many other bacteria of industrial importance.

As we have shown throughout this work, for example by use of the eGFP ORF, the compatibility of EcoFlex and BacilloFlex greatly benefits their utility. Widespread adoption of the EcoFlex standard, and its derivatives like BacilloFlex, will expand the range of available tools and facilitate shuttling of parts between organisms. We hope that both academic and industrial researchers will benefit from the utility of BacilloFlex, and that this work will form part of a standardized modular cloning scheme contributing to realizing the synthetic biology potential of *B. subtilis* and related gram-positive bacteria.

## 4. Materials and Methods

### Strains and culture conditions

Routine growth of *Escherichia coli* K12 JM109 (NEB, gift from Marcos Valenzuela) and *Bacillus subtilis* 168 was carried out in Luria-Bertani (LB) agar or broth at 37°C (shaking at 200 rotations per minute, RPM, for liquid culture). *E. coli* was selected using carbenicillin (100 μg/ml, Fisher Scientific) or chloramphenicol (40 μg/ml, Acros Organics)(cml40) and *B. subtilis* with 10 μg/ml chloramphenicol (cml10).

#### *E. coli* transformation

Chemically competent *E. coli* cells were prepared as described by Chung et al. (1989). To transform, cells were incubated with DNA for 30 min on ice, heat-shocked at 42°C for 90 s and returned to ice for 5 min. 0.9 ml LB-broth was added and cells allowed to recover at 37°C, 200 RPM for 90 min before plating onto selective media.

#### *B. subtilis* transformation

Preparation of *B. subtilis* competent cells, storage and transformation were performed using the Groningen method as described by Bron (1990). Aliquots were frozen and stored at -80°C. 200 μl of transformation culture was plated onto LB + cml10, and the remaining 800 μl added to 5 ml LB with a final concentration of 5 μg/ml chloramphenicol. This culture was grown overnight at 37°C 200 RPM to allow outgrowth of transformants which could be plated subsequently if no colonies were obtained on the initial plate.

### Routine molecular biology

All oligonucleotide sequences are detailed in the supplementary information (Table S1). Routine isolation of plasmid DNA was performed using the QIAprep Miniprep kit (Qiagen) according to the manufacturer’s instructions. DNA concentrations were measured using a NanoDrop 2000 (Thermo Scientific). Routine verification of assembly or chromosomal integration was performed by colony PCR using *Taq* DNA polymerase (NEB). Where possible functional assays, e.g. GFP expression, were used to screen for correct assembly. DNA in agarose gels was visualized using SafeView nucleic acid stain (NBS Biologicals).

### Part construction

#### Constructing parts by oligo annealing

Promoters, RBS, NTTs (Epr, PhrK), XC5 and XC3 were ordered as oligonucleotides (Sigma-Aldrich) with relevant flanking BsaI sites and overhangs for cloning into pBP-lacZ as described by Moore et al. (2016). Equimolar amounts (10 μM) of complimentary oligos with upstream 5’ NdeI and downstream 3’ SphI compatible overhangs were combined in a PCR tube, heated to 95°C for 10 s and then cooled to 10°C at 0.2°C s^-1^. Annealed oligos were phosphorylated by incubation with 10 units of T4 Polynucleotide Kinase in 1X T4 DNA ligase buffer (NEB) for 1 hour at 37°C followed by 20 min at 65°C. Parts exceeding 120 bp were designed as two pairs of oligonucleotides with compatible 4 bp overhangs such that they would ligate to form the full-length part with upstream 5’ NdeI and downstream 3’ SphI overhangs.

#### Constructing parts by PCR

NTTs (cotB, cotY) and HAs were amplified by PCR from a colony of *B. subtilis* 168 resuspended in 50 μl nuclease-free water (Qiagen) using Q5® High-Fidelity DNA Polymerase (NEB). SA was amplified from a gBlocks^®^ SA template (Integrated DNA technologies). XylE, cat and catXC were amplified from the Bacillosensor plasmid. Primers included 5’ overhangs with relevant restriction and fusion sites. After size verification by gel electrophoresis, the remaining PCR reaction volume was purified using the QIAquick PCR purification kit (Qiagen). For level 1 parts approximately 500 ng of purified PCR product was digested using 20 units of BsaI-HF in 50 μl 1X NEBuffer 3.1 (NEB) for 1 hour at 37°C followed by 20 min at 65°C. Level 0 parts digested as level 1 but using 20 units of each NdeI and SphI-HF (NEB) in 1X CutSmart® buffer (NEB).

#### Cloning parts into plasmids

For cloning, pBP and pTU1 vectors were digested with NdeI/SphI and BsaI-HF respectively, using the conditions described for their cognate parts above, but using 1 μg of DNA. The desired backbone portion was purified using gel electrophoresis and the QIAquick Gel Extraction kit (Qiagen). 100 ng of digested insert was mixed with 10 ng of digested backbone vector in 20 μl 1X T4 DNA ligase buffer (NEB) and 400 units T4 DNA ligase (NEB) and incubated for 30-60 minutes at room temperature. Blue/white screening for insertion of bio-parts into pBP-lacZ was performed on LB + cml40 supplemented with 0.1 mM isopropyl β-D-1-thiogalactopyranoside (IPTG, Melford Laboratories Ltd) and 40 μg/ml 5-bromo-4-chloro-3-indolyl-β-D-galactopyranoside (X-gal, Melford Laboratories Ltd). All parts were stored in *E. coli* K12 JM109 (NEB).

### Sequencing

Level 0 bio-parts and level 1 homology arms were completely sequenced using the BigDye® Terminator v3.1 Cycle Sequencing Kit (AppliedBiosystems™) according to the manufacturer’s instructions. Samples were sent to Edinburgh Genomics (https://genomics.ed.ac.uk/) for analysis.

### Level 1 Golden Gate Reaction

40 fmol and 20 fmol per 20 μl reaction of each part and backbone vector respectively were combined in 1X T4 DNA ligase buffer containing 20 units of BsaI-HF (NEB) and 100 units T4 DNA ligase. The reaction was cycled 25 times for 90 s at 37°C and 3 min at 16°C. Following this cycle, the reaction was held at 50°C and then 80°C for 5 min each. Routinely 20 μl reactions were transformed into *E. coli*.

### Level 2 Golden Gate Reaction

Reaction conditions were as for level 1 Golden Gate Reactions but containing 20 units of BsmBI (NEB) in place of BsaI-HF. Reactions were cycled 25 times for 3 min at 42°C and 3 min at 16°C. This was followed by 10 min incubation at 80°C. Routinely 5-20 and/or 15-20 μl were transformed into *E. coli* and *B. subtilis* respectively.

### Fluorescence measurements

Single colonies of *B. subtilis* strains were picked and grown overnight in LB (+ cml10 as appropriate) at 30°C. Cell density was measured at 600 nm using a FLUOstar Omega (BMG Labtech) plate reader. Strains were subcultured into 100 μl LB (+ cml10 as appropriate) to a starting OD_600_ of ∼0.005 in triplicate in a randomized layout in a black F-bottom 96-well plate (Greiner Bio-One) and sealed with AeraSeal™ (Porvair Sciences). Arsenate was added from a 10000 ppb or 10 μg/ml arsenate (22.4 μg/ml sodium arsenate heptahydrate, Sigma-Aldrich, in sterile water) stock solution as needed. Cells were grown for 16 hours at 30°C 600 RPM in a FLUOstar Omega (BMG Labtech) recording OD_600_ and eGFP fluorescence (excitation 485 nm, emission 520nm), blanked against LB, every 10 min.

### Promoter activity data analysis

Constitutive promoter activity was quantified during exponential phase (OD_600_ ∼ 0.3 – 0.6). Average relative fluorescence units (RFU) of triplicate *B. subtilis* wild-type cultures was subtracted from RFU measurements of strains. To correct for cell density RFUs were divided by OD_600_. RFU/OD_600_ triplicate data were averaged over a 50 min period (6 time points).

P*_ars_* activity was quantified during stationary phase. RFU/OD_600_ triplicate data (wild-type fluorescence not subtracted) were averaged over a 1 hour period (7 time points).

## 5. Acknowledgements

We thank Dr Chao-Kuo Liu and Marcos Valenzuela for useful discussion.

## References

Amuguni H. & Tzipori S. 2012. *Bacillus subtilis* A temperature resistant and needle free delivery system of immunogens. Human Vaccines & Immunotherapeutics, 8, 8.

Bloor A. E. & Cranenburgh R. M. 2006. An efficient method of selectable marker gene excision by Xer recombination for gene replacement in bacterial chromosomes. Applied and Environmental Microbiology, 72, 2520–2525.

Brockmeier, U., Caspers, M., Freudl, R., Jockwer, A., Noll, T. & Eggert, T. 2006. Systematic screening of all signal peptides from *Bacillus subtilis*: A powerful strategy in optimizing heterologous protein secretion in gram-positive bacteria. Journal of Molecular Biology, 362, 393–402.

Bron S. 1990. Plasmids. In: Molecular Biological Methods for Bacillus (Modern Microbiological Methods) Harwood, C. R. & Cutting, S. M. (eds.) Wiley & Sons.

Carrasco, B., Yadav, T., Serrano, E. & Alonso, J. C. 2015. *Bacillus subtilis* Reco and Ssba are crucial for Reca-mediated recombinational Dna repair. Nucleic Acids Research, 43, 5984–5997.

Chen, X. T., Ji, J. B., Liu, Y. C., Ye, B., Zhou, C. Y. & Yan, X. 2016. Artificial induction of genetic competence in *Bacillus amyloliquefaciens* isolates. Biotechnology Letters, 38, 2109–2117.

Chung, C. T., Niemela, S. L. & Miller, R. H. 1989. One-step preparation of competent *Escherichia coli* - transformation and storage of bacterial cells in the same solution. Proceedings of the National Academy of Sciences of the United States of America, 86, 21722175.

Cutting, S. M., Hong, H. A., Baccigalupi, L. & Ricca, E. 2009. Oral vaccine delivery by recombinant spore probiotics. International Reviews of Immunology, 28, 487–505.

De SOUZA R. D., Batista, M. T., Luiz, W. B., Cavalcante, R. C. M., Amorim, J. H., Bizerra, R. S. P., Martins, E. G. & Ferreira, L. C. D. 2014. *Bacillus subtilis* spores as vaccine adjuvants: further insights into the mechanisms of action. Plos One, 9, 10.

Dubnau D. 1991. Genetic competence in *Bacillus subtilis*. Microbiological Reviews, 55, 395–424.

Engler, C., Kandzia, R. & Marillonnet, S. 2008. A one pot, one step, precision cloning method with high throughput capability. Plos One, 3, 7.

Flanagan, S. V., Johnston, R. B. & Zheng, Y. 2012. Arsenic in tube well water in Bangladesh: health and economic impacts and implications for arsenic mitigation. Bulletin of the World Health Organization, 90, 839–846.

Fukushima, T., Ishikawa, S., Yamamoto, H., Ogasawara, N. & Sekiguchi, J. 2003. Transcriptional, functional and cytochemical analyses of the *veg* gene in *Bacillus subtilis*. Journal of Biochemistry, 133, 475–483.

Guiziou, S., Sauveplane, V., Chang, H. J., Clerte, C., Declerck, N., Jules, M. & Bonnet, J. 2016. A part toolbox to tune genetic expression in *Bacillus subtilis*. Nucleic Acids Research, 44, 7495–7508.

Hao, T., Han, B. B., Ma, H. W., Fu, J., Wang, H., Wang, Z. W., Tang, B. C., Chen, T. & Zhao, X. M. 2013. *In silico* metabolic engineering of *Bacillus subtilis* for improved production of riboflavin, Egl-237, (R,R)-2,3-butanediol and isobutanol. Molecular Biosystems, 9, 2034–2044.

Harwood C. R. & Cranenburgh R. 2008. *Bacillus* protein secretion: an unfolding story. Trends in Microbiology, 16, 73–79.

Henkin, T. M., Grundy, F. J., Nicholson, W. L. & Chambliss, G. H. 1991. Catabolite repression of alpha-amylase gene-expression in *Bacillus subtilis* involves a trans-acting gene product homologous to the *Escherichia coli Laci* and *Galr* repressors. Molecular Microbiology, 5, 575–584.

Jordan, S., Junker, A., Helmann, J. D. & Mascher, T. 2006. Regulation of Liars-dependent gene expression in *Bacillus subtilis*: Identification of inhibitor proteins, regulator binding sites, and target genes of a conserved cell envelope stress-sensing two-component system. Journal of Bacteriology, 188, 5153–5166.

Knurr, J., Benedek, O., Heslop, J., Vinson, R. B., Boydston, J. A., Mcandrew, J., Kearney, J. F. & Turnbough, C. L. 2003. Peptide ligands that bind selectively to spores of *Bacillus subtilis* and closely related species. Applied and Environmental Microbiology, 69, 6841–6847.

Kunst, F., Ogasawara, N., Moszer, I., Albertini, A. M., Alloni, G., Azevedo, V., Bertero, M. G., Bessieres, P., Bolotin, A., Borchert, S., Borriss, R., Boursier, L., Brans, A., Braun, M., Brignell, S. C., Bron, S., Brouillet, S., Bruschi, C. V., Caldwell, B., Capuano, V., Carter, N. M., Choi, S. K., Codani, J. J., Connerton, I. F., Cummings, N. J., Daniel, R. A., Denizot, F., Devine, K. M., Dusterhoft, A., Ehrlich, S. D., Emmerson, P. T., Entian, K. D., Errington, J., Fabret, C., Ferrari, E., Foulger, D., Fritz, C., Fujita, M., Fujita, Y., Fuma, S., Galizzi, A., Galleron, N., Ghim, S. Y., Glaser, P., Goffeau, A., Golightly, E. J., Grandi, G., Guiseppi, G., Guy, B. J., Haga, K., Haiech, J., Harwood, C. R., Henaut, A., Hilbert, H., Holsappel, S., Hosono, S., Hullo, M. F., Itaya, M., Jones, L., Joris, B., Karamata, D., Kasahara, Y., Klaerrblanchard, M., Klein, C., Kobayashi, Y., Koetter, P., Koningstein, G., Krogh, S., Kumano, M., Kurita, K., Lapidus, A., Lardinois, S., Lauber, J., Lazarevic, V., Lee, S. M., Levine, A., Liu, H., Masuda, S., Mauel, C., Medigue, C., Medina, N., Mellado, R. P., Mizuno, M., Moestl, D., Nakai, S., Noback, M., Noone, D., Oreilly, M., Ogawa, K., Ogiwara, A., Oudega, B., Park, S. H., Parro, V., Pohl, T. M., Portetelle, D., Porwollik, S., Prescott, A. M., Presecan, E., Pujic, P., Purnelle, B., et al. 1997. The complete genome sequence of the Gram-positive bacterium *Bacillus subtilis*. Nature, 390, 249–256.

Lee, S. Y., Park, J. H., Jang, S. H., Nielsen, L. K., Kim, J. & Jung, K. S. 2008. Fermentative butanol production by clostridia. Biotechnology and Bioengineering, 101, 209–228.

Liu, W. X., Fu, J., Wang, Z. W. & Chen, T. 2014. Elimination of carbon catabolite repression in *Bacillus subtilis* for the improvement of 2,3-butanediol production. Proceedings of the 2012 International Conference on Applied Biotechnology (Icab 2012), Vol 1, 249, 323–331.

Mock M. & Fouet A. 2001. Anthrax. Annual Review of Microbiology, 55, 647–671.

Moore, S. J., Lai, H. E., Kelwick, R. J. R., Ghee, S. M., Bell, D. J., Polizzi, K. M. & Freemont, P. S. 2016. Ecoflex: A multifunctional Moclo kit for *E. coli* synthetic biology. Acs Synthetic Biology, 5, 1059–1069.

Narula, J., Fujita, M. & Igoshin, O. 2016. Functional requirements of cellular differentiation: lessons from *Bacillus subtilis*. Current Opinion in Microbiology, 34, 38–46.

Pohl, S., Bhavsar, G., Hulme, J., Bloor, A. E., Misirli, G., Leckenby, M. W., Radford, D. S., Smith, W., Wipat, A., Williamson, E. D., Harwood, C. R. & Cranenburgh, R. M. 2013. Proteomic analysis of *Bacillus subtilis* strains engineered for improved production of heterologous proteins. Proteomics, 13, 3298–3308.

Polissi A. & Harayama S. 1993. *In vivo* reactivation of catechol 2,3-dioxygenase mediatd by a chloroplast-type ferredoxin - a bacterial strategy toexpand the substrate specificity of aromatic degradative pathways. Embo Journal, 12, 3339–3347.

Quentin, Y., Fichant, G. & Denizot, F. 1999. Inventory, assembly and analysis of *Bacillus subtilis* Abc transport systems. Journal of Molecular Biology, 287, 467–484.

Radeck, J., Kraft, K., Bartels, J., Cikovic, T., Durr, F., Emenegger, J., Kelterborn, S., Sauer, C., Fritz, G., Gebhard, S. & Mascher, T. 2013. The Bacillus Biobrick Box: generation and evaluation of essential genetic building blocks for standardized work with Bacillus subtilis. Journal of Biological Engineering, 7, 16.

Rebatchouk, D., Daraselia, N. & Narita, J. O. 1996. Nomad: A versatile strategy for in vitro Dna manipulation applied to promoter analysis and vector design. Proceedings of the National Academy of Sciences of the United States of America, 93, 10891–10896.

Sato T. & Kobayashi Y. 1998. The *ars* operon in the skin element of *Bacillus subtilis* confers resistance to arsenate and arsenite. Journal of Bacteriology, 180, 1655–1661.

Sauer C., Syvertsson, S., Bohorquez, L. C., Cruz, R., Harwood, C. R., Van RIJ, T. & Hamoen L. W. 2016. Effect of genome position on heterologous gene expression in *Bacillus subtilis*: an unbiased analysis. Acs Synthetic Biology, 5, 942–947.

Schallmey, M., Singh, A. & Ward, O. P. 2004. Developments in the use of *Bacillus* species for industrial production. Canadian Journal of Microbiology, 50, 1–17.

Spizizen J. 1958. Transformation of biochemically deficient strains of *Bacillus subtilis* by deoxyribonucleate. Proceedings of the National Academy of Sciences of the United States of America, 44, 1072–1078.

Sun, H. G., Bie, X. M., Lu, F. X., Lu, Y. P., Wu, Y. D. L. & Lu, Z. X. 2009. Enhancement of surfactin production of *Bacillus subtilis Fmbr* by replacement of the native promoter with the Pspac promoter. Canadian Journal of Microbiology, 55, 1003–1006.

Van DIJL, J. M. & Hecker, M. 2013. *Bacillus subtilis*: from soil bacterium to super-secreting cell factory. Microbial Cell Factories, 12, 6.

Wang, H., Wang, Y. X. & Yang, R. J. 2017. Recent progress in *Bacillus subtilis* spore-surface display: concept, progress, and future. Applied Microbiology and Biotechnology, 101, 933949.

Wu, X. C., Lee, W., Tran, L. & Wong, S. L. 1991. Engineering a *Bacillus subtilis* expression-secretion system with a strain deficient in 6 extracellular proteases. Journal of Bacteriology, 173, 4952–4958.

Yadav, T., Carrasco, B., Serrano, E. & Alonso, J. C. 2014. Roles of *Bacillus subtilis* Dpra and Ssba in Reca-mediated Genetic Recombination. Journal of Biological Chemistry, 289, 27640–27652.

Zhou, Z. W., Dong, H., Huang, Y. M., Yao, S. W., Liang, B. S., Xie, Y. Q., Long, Y., Mai, J. L. & Gong, S. T. 2017. Recombinant *Bacillus subtilis* spores expressing cholera toxin B subunit and *Helicobacter pylori* urease B confer protection against *H. pylori* in mice. Journal of Medical Microbiology, 66, 83–89.

